# JQ1 affects BRD2-dependent and independent transcription regulation without disrupting H4-hyperacetylated chromatin states

**DOI:** 10.1101/215442

**Authors:** Lusy Handoko, Bogumil Kaczkowski, Chung-Chau Hon, Marina Lizio, Masatoshi Wakamori, Takayoshi Matsuda, Takuhiro Ito, Prashanti Jeyamohan, Yuko Sato, Kensaku Sakamoto, Shigeyuki Yokoyama, Hiroshi Kimura, Aki Minoda, Takashi Umehara

## Abstract

The bromodomain and extra-terminal domain (BET) proteins are promising drug targets for cancer and immune diseases. However, BET inhibition effects have been studied more in the context of bromodomain-containing protein 4 (BRD4) than BRD2, and the BET protein association to histone H4-hyperacetylated chromatin is not understood at the genome-wide level. Here, we report transcription start site (TSS)-resolution integrative analyses of ChIP-seq and transcriptome profiles in human non-small cell lung cancer (NSCLC) cell line H23. We show that di-acetylation at K5 and K8 of histone H4 (H4K5acK8ac) co-localizes with H3K27ac and BRD2 in the majority of active enhancers and promoters, where BRD2 has a stronger association with H4K5acK8ac than H3K27ac. Interestingly, although BET inhibition by JQ1 led to complete reduction of BRD2 binding to chromatin, only local changes of H4K5acK8ac levels were observed. In addition, a remarkable number of BRD2-bound genes, including *MYC* and its downstream target genes, were transcriptionally upregulated upon JQ1 treatment. Using BRD2-enriched sites and transcriptional activity analysis, we identified candidate transcription factors potentially involved in the JQ1 response in BRD2-dependent and independent manner.

---

Lysine acetylation of core histones is one of the major post-translational modifications involved in the control of eukaryotic gene expression^1^. Acetylated status of lysine residue is mainly recognized by bromodomains which are found in many chromatin-associated proteins^2^. Mammalian bromodomain and extra-terminal domain (BET) family comprising BRD2, BRD3, BRD4 and BRDT is involved in transcriptional regulation of cancer-related genes and has been recognized as a therapeutic target in many cancer types, such as NUT midline carcinoma (NMC)^3^ and acute myeloid leukemia (AML)^4^. JQ1, an inhibitor of BET proteins that binds the acetyllysine (Kac)-binding pocket of bromodomain^3^, has been shown to selectively suppress the expression of tumor-promoting genes in cancer cells and emerges as a promising anti-cancer therapeutic drug^3,5–9^. Since studies on BET inhibition mechanisms have mostly focused on the correlation between BRD4 genomic occupancy and JQ1-induced transcriptional changes^5,10^, little is known on how BRD2 and hyperacetylated histone H4, as the direct chromatin target of BET proteins, engage in the cancer-related pathways.

The BET family shares a C-terminal extra-terminal (ET) domain and two tandem bromodomains, BD1 and BD2 that primarily bind to multi-acetylated histone H4 tail at K5, K8, K12, and K16 *in vitro*, but not to the mono-acetylated histone H3/H4 peptides, including that of acetylated H3 K27 (H3K27ac)^8,9,11,12^. Since acetylation of histone H4 in the nucleus is proposed to occur at K16 first, then at K12, K8, and K5^13^, the simultaneous acetylation of K5 and K8 indicates a typical state of histone H4 hyperacetylation^13^. Indeed, BET proteins preferentially bind to the K5/K8-diacetylated H4 tail peptides mimicking hyperacetylated H4 *in vitro*^11,12^. Additionally, H4 K5 acetylation (H4K5ac) facilitated by EP300 (histone acetyltransferase p300) and disruptor of telomere silencing 1-like (DOT1L) might facilitate the binding of BRD4 to the chromatin^14^.

At a genome-wide level, BRD2, BRD3, and BRD4 co-occupy H3K27ac sites at most of the active promoters together with RNA polymerase II and Mediator complex^15,16^. However, it is unclear whether BET proteins have a preferred binding to hyperacetylated histone H4 or H3K27ac *in vivo*. Furthermore, while genome-wide H3K27ac-enriched sites that overlap highly with BRD2, BRD3, and BRD4 sites are not sensitive to a certain concentration of JQ1^5,17^, effects of JQ1 to hyperacetylated histone H4 are not studied at the genome-wide level. The lack of antibody specifically recognizing the hyperacetylated states of histone H4 tail potentially hampers efforts to address this question.

In this study, we generated a monoclonal antibody that specifically recognizes simultaneous acetylations at K5 and K8 of histone H4 (referred to as H4K5acK8ac), and employed it for ChIP-seq analyses. We used non-small cell lung cancer (NSCLC) cell line H23, which carries *KRAS* and *TP53* mutations and has a mild sensitivity to BET inhibitors JQ1 and iBET^10,18,19^, to assess the co-occupancy of BRD2 with hyperacetylated H4 and H3K27ac at a genome-wide level. Furthermore, we performed transcriptomics by cap-analysis of gene expression (CAGE) method^20^. CAGE captures 5’ starts of transcribed RNAs, making it possible to determine precise positions of TSSs and transcribed enhancers genome-wide^20^. Through the integrative analyses of ChIP-seq and CAGE data, we elucidate the involvement of BRD2 in gene regulation upon BET inhibition by JQ1 in H23 cells.

## RESULTS

### Generation of a specific monoclonal antibody against H4K5acK8ac

In order to explore the association of BRD2 with histone H4 hyperacetylation at a genome-wide level and to elucidate the effect of JQ1 on hyperacetylated H4, we first developed a mouse monoclonal antibody that specifically recognizes hyperacetylated histone H4. We synthesized the full-length histone H4 protein with site-specific acetylations at K5 and K8 on a milligram scale^21^, and used it as the antigen. Through screening with ELISA and Western blot, we successfully obtained two hits of hybridoma (1A9D7 and 2A7D9) that recognize the histone H4 proteins containing di-acetylation of K5acK8ac (*i.e.* H4K5acK8ac and tetra-acetylated H4) but not mono-acetylated H4 proteins (**Fig. 1a** and **Supplementary Fig. 1**). These were further confirmed by the peptide array analysis showing that 2A7D9 binds to H4 peptide only if K5acK8ac is present (**Fig. 1b** and **Supplementary Table 1**), whereas 1A9D7 also weakly bound to H2B K12acK15ac (**Supplementary Table 1**). For comparison, we also examined specificity of the two commercially available antibodies for hyperacetylated H4 (*i.e*. Upstate 06-946 and Abcam ab177790), which we found not to be specific (**Supplementary Table 1**). In addition to tetra-acetylated histone H4 or H4K5acK8ac, these antibodies also detected H4 mono-acetylation (K5 and K12) (**Supplementary Table 1**).

**Figure 1.**
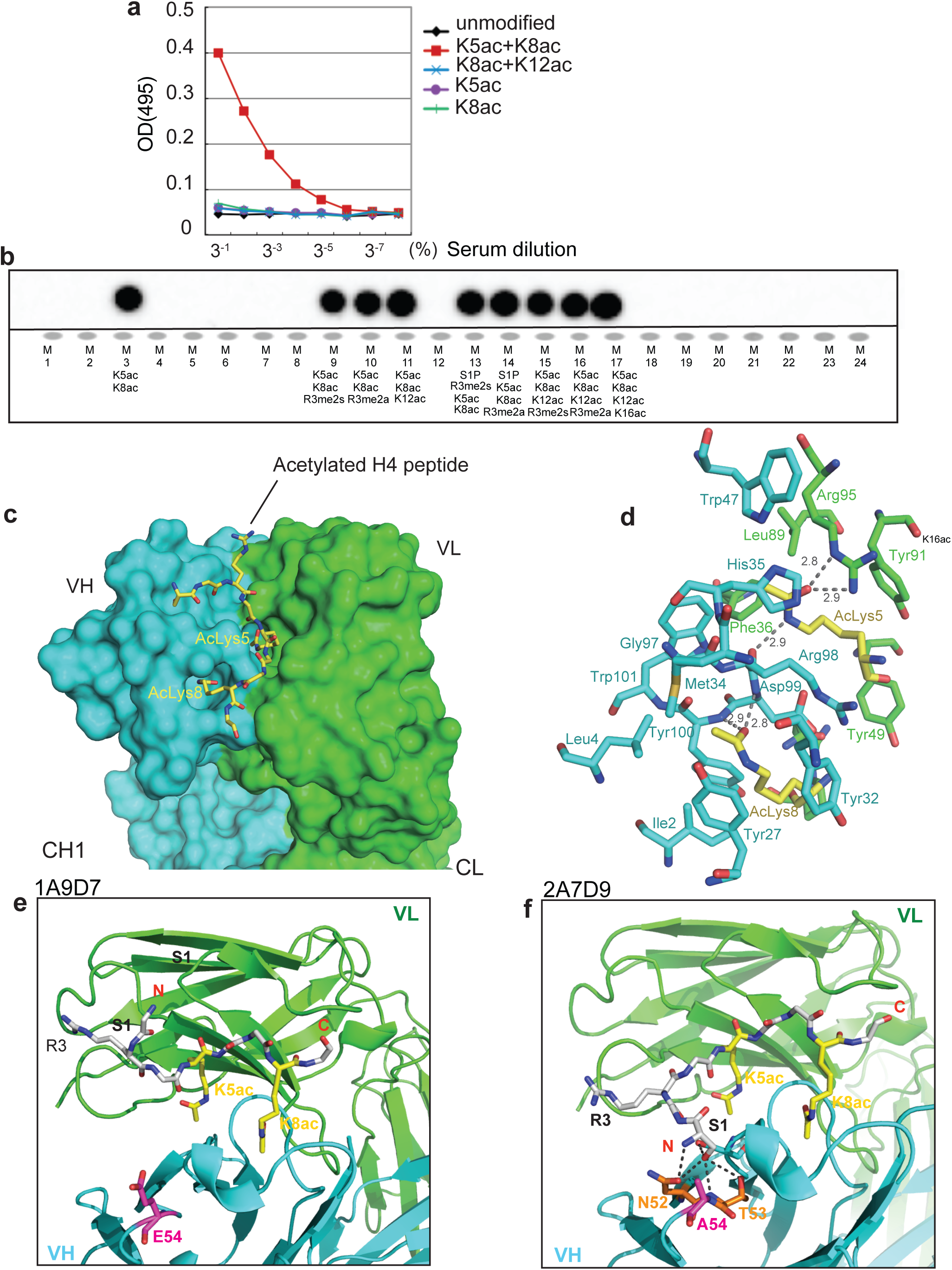
Development of monoclonal antibodies that specifically recognize H4K5acK8ac. **(a)** ELISA assay for 2A7D9 antibody. Histone H4 protein containing indicated site-specific acetyllysine(s) are used as antigens. **(b)** Specificity analysis of 2A7D9 antibody using modified histone peptide array. Histone H4 peptides containing both K5ac and K8ac are indicated in the bottom panel. **(c)** Crystal structure of the 2A7D9 Fab fragment in complex with K5/K8-diacetylated histone H4 (1-12) peptide. Heavy and light chains are depicted in cyan and green, respectively. The K5/K8-diacetylated H4 peptide is shown in a yellow stick. **(d)** Close-up view of the 2A7D9 Fab fragment in complex with the H4 peptide. Carbon atoms of VH and VL chains are shown in cyan and green, respectively. Carbon atoms of acetyllysines positioned at 5 and 8 in the K5/K8-diacetylated H4 peptide are shown in yellow. All the nitrogen and oxygen atoms are shown in blue and red, respectively. Hydrogen bonds are indicated as dotted lines with linear distances (Å). **(e and f)** Close-up view of the complex around the 54th residue of VH chain. **(e)** 1A9D7antibody, **(f)** 2A7D9 antibody. In **(e)** and **(f)**, N- and C-terminus of the K5/K8-diacetylated H4 peptide is shown as red label of N and C, respectively. Glutamic acid at 54th position **(e)** and Alanine at 54th position **(f)** is shown in magenta. In **(f)**, asparagine 52 and threonine 53 are shown in orange, and hydrogen bonds formed between the VH chain (i.e. N52–A54) and the H4 peptide (i.e. S1) are shown as dotted lines.

We next determined crystal structures of the antigen-binding (Fab) fragments of the two monoclonal antibodies, 2A7D9 and 1A9D7, in a complex with the H4K5acK8ac tail peptide at 1.7–1.8 Å. With both antibodies, the H4K5acK8ac peptide is recognized by a cleft formed by V_H_ and V_L_, and the K5ac and K8ac side chains are respectively recognized by a different cavity (**Supplementary Table 2** and **Fig. 1c**). The carbonyl group, which is generated by acetylation of the ε-amino group of K5ac, is specifically recognized by the side-chain guanidium group of R95 in V_L_ and the main-chain carbonyl group of R98 in V_H_. On the other hand, the carbonyl group of the ε-amino group of K8ac is specifically recognized by the main-chain amino groups of D99 and Y100 in V_H_ (**Fig. 1d**). The side chain of R98 spatially separates the two Kac-recognizing cavities, and only V_H_ residues form the K8ac-recognizing cavity (**Fig. 1d**). Hence, we found that the individual H4K5ac- and H4K8ac-binding modes of the antibodies are completely different from the cooperative H4K5acK8ac-binding mode of the BRDT BD1 bromodomain reported by Moriniere *et al*.^22^

By comparing the primary and tertiary structures of the two antibodies (**Fig. 1e,f** and **Supplementary Fig. 1c**), we reasoned that A54 of V_H_ in 2A7D9 is responsible for the strict discrimination of H4K5acK8ac. In 1A9D7 composing E54 at the same position, the Fab fragment does not bind to the N-terminal S1 residue of H4 (**Fig. 1e**). In contrast, A54 of 2A7D9 has a smaller side chain than that of the glutamic acid (*i.e*. E54), thus allowing binding of N52 and T53 to S1 of the H4 N-terminal tail (**Fig. 1f**). Thus, 2A7D9 preferentially binds to the N-terminal H4 sequence that contains S1 at the −4 position relative to H4K5ac than to any other histone tail peptide sequences (**Fig. 1b**). Since 1A9D7 cannot bind to H4S1, it cannot exclude binding to the N-terminal H2B sequence carrying K12ac and K15ac that contains P8 at the −4 position relative to H2B K12ac (**Supplementary Fig. 1c**). Although there are a series of monoclonal antibodies that detect mono-acetylation of histone H4 at K5, K8, K12 and K16, no structurally validated monoclonal antibody recognizing a specific combination of histone tail modifications has been reported before to our knowledge. Hence, our crystallographic analysis provides the structural basis for the selective recognition and validation of H4K5acK8ac by the 2A7D9 antibody.

### H4K5acK8ac co-localizes with H3K27ac at active promoters and enhancers

To perform a genome-wide profiling of H4 hyperacetylation and to compare it with other active chromatin marks, we performed ChIP-seq with the 2A7D9 antibody against H4K5acK8ac, along with antibodies for H3K27ac, H3K4me1, and H3K4me3. We identified 22,882 robust H4K5acK8ac peaks (irreproducible discovery rate, IDR < 0.01), of which 34.5% are at the RefSeq promoters, 25.6% at introns and 29.8% at intergenic regions (**Fig. 2a** and **Supplementary Table 3**). Consistent with the notion of acetylated histone H4 as an open chromatin mark, H4K5acK8ac is found at transcriptionally active or open chromatin regions, including: i) enhancers (overlap with H3K27ac and H3K4me1; **Fig. 2b,c**), ii) active enhancers (H3K27ac >5 kb outside TSS; **Fig. 2d**, top), iii) active promoters (H3K4me3 5 kb ± from TSS; **Fig. 2d**, bottom), and iv) FANTOM5 transcribed enhancers (**Fig. 2e**). The co-localization of H4K5acK8ac with the active promoter and enhancer marks were further confirmed by immunostaining (**Supplementary Fig. 2a**). Consistently, H4K5acK8ac is associated with actively transcribed promoters and enhancers, as determined by co-localization with CAGE signals (~77% of TSS with expression level ≥ 0.25 CAGE tag per million, TPM) (**Supplementary Table 3**). Overall, our data show that the H4K5acK8ac mostly occupies active enhancer and promoter regions.

**Figure 2.**
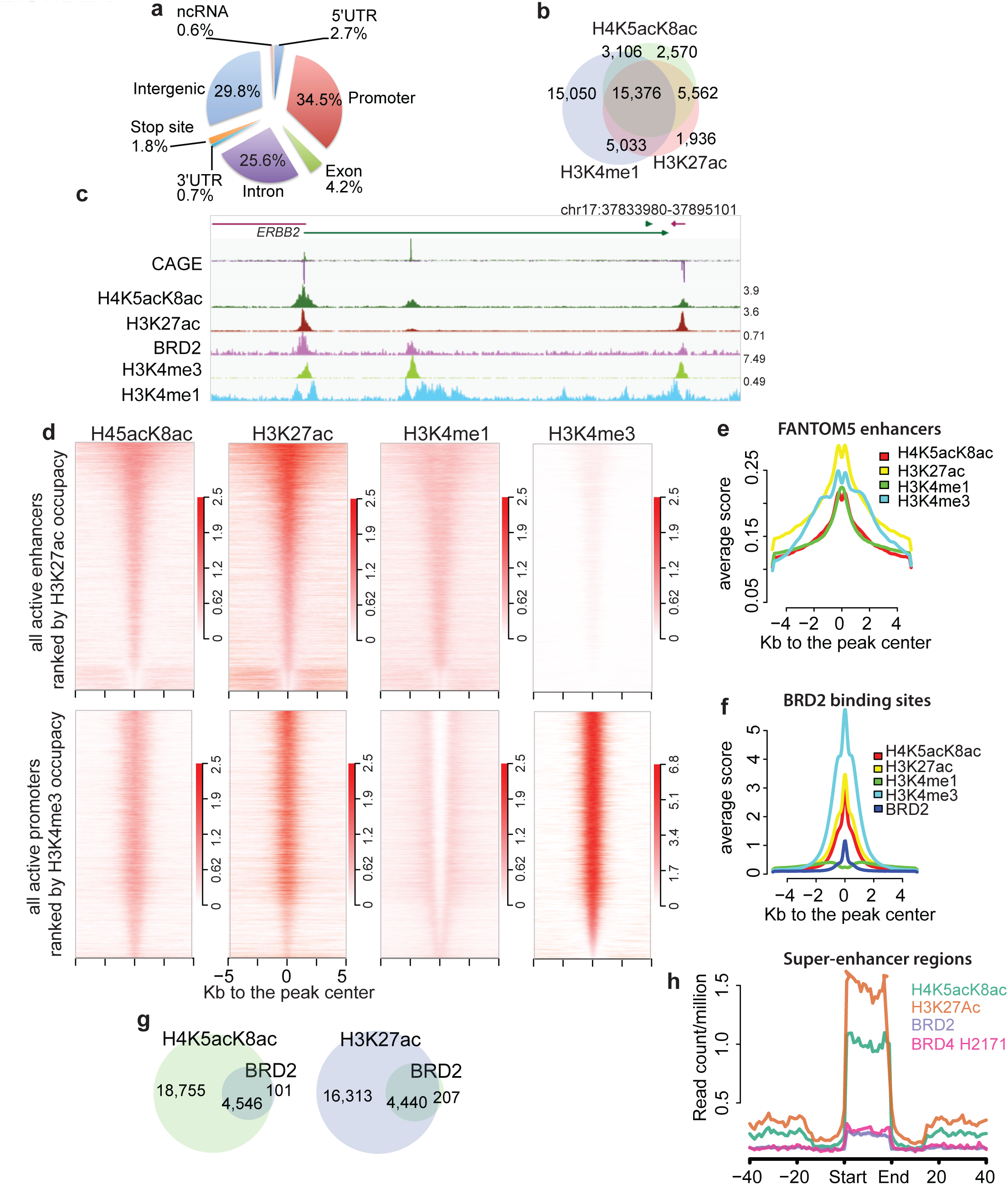
Genome-wide characterization of H4K5acK8ac sites and BRD2 binding sites. **(a)** A chart of genomic localization of H4K5acK8ac peaks. **(b)** Venn diagram showing a high overlap of H4K5acK8ac with two enhancer marks H3K27ac and H3K4me1. Using a simple intersection with at least 1 bp overlap, 78 % of H4K5acK8ac sites overlap with 87% of H3K27ac sites and 80% of H4K5acK8ac with 50% of H3K4me1. **(c)** An example of a genome browser view of ChIP-seq signal of H4K5acK8ac, BRD2, other histone marks and gene expression signals by CAGE. **(d)** Heatmap of normalized H4K5acK8ac ChIP-seq intensities (read per million, rpm) within 5 kb +/-from the summit of active enhancers defined by H3K27ac located outside promoter (top) and active promoter defined by H3K4me3 at the TSS (bottom). H3K4me1 and H3K27ac mark active enhancer (top), while active promoters are highly enriched for H3K4me3 and H3K27ac but devoid of H3K4me1 (bottom). Color density indicates the enrichments of histone marks. **(e)** Enrichment of H4K5acK8ac and other histone marks within FANTOM5-defined enhancer regions. Average profiles (average reads per million) of histone marks were plotted within 5 kb +/-from the center of the enhancers (plotted by ngs.plot package41). **(f)** Enrichment of active histone modifications within BRD2 binding sites, as shown by average ChIP-seq profiles of the histone marks. **(g)** Venn diagram showing a high overlap of BRD2-H4K5acK8ac (97.8%, left) and BRD2-H3K27ac (95.5%, right). **(h)** Enrichment of BRD2 and BRD4 ChIP-seq intensities (average rpm) in the super-enhancer regions identified by H4K5acK8ac ChIP-seq. BRD4 ChIP-seq from SCLC H2171 cell line was obtained from GSE4235513.

### Stronger association between BRD2 and H4K5acK8ac than BRD2 and H3K27ac

Although BET proteins preferentially bind to multi-acetylated histone H4 *in vitro* (mimicking hyperacetylation), it is not known whether BRD2 preferentially binds to hyperacetylated H4 *in vivo*, compared to other acetylated forms such as H3K27ac. Since the BRD2 binding pattern in H23 cells has not been characterized yet, we performed BRD2 ChIP-seq and identified 4,647 robust BRD2 binding sites, which are found at promoters (47.5%), introns (21.7%), and at intergenic regions (15.8%) (**Supplementary Fig. 2b**). Moreover, when overlaid with the CAGE data, 87% of BRD2 sites are localized at actively transcribed promoters of coding and non-coding genes (CAGE ≥ 0.25 TPM, **Supplementary Table 3**). Consistently, BRD2 sites are highly enriched for active marks (*e.g.*, H3K4me3, H3K27ac, and H4K5acK8ac) (**Fig. 2f**), where at least 95% of the BRD2 peaks overlap with H4K5acK8ac and H3K27ac (**Fig. 2g**).

In a complementary analysis, we examined the enrichment of bromodomain proteins BRD2, BRD4 and EP300 at the H4K5acK8ac sites using the publicly available ChIP-seq datasets from the closely related cell types: BRD4 ChIP-seq from SCLC cell line H2171 (GSM1038270)^17^ and ENCODE P300 ChIP-seq from NSCLC A549^23^. We found that promoter- and enhancer-associated H4K5acK8ac regions were enriched for BRD2, BRD4 and EP300 (**Supplementary Fig. 2c**), suggesting the co-localization of H4K5acK8ac with these bromodomain-containing proteins. We also identified 432 super-enhancers based on H4K5acK8ac peaks, which are also enriched with H3K27ac and BRD2 (**Fig. 2h** and **Supplementary Table 4**). Taken together, our ChIP-seq analysis shows that BRD2 mostly localizes to active promoters and enhancers including super-enhancers in H23 cells.

To determine how strongly BRD2 co-occurs with H4K5acK8ac or H3K27ac, we calculated the odds ratio (OR) between BRD2 and each histone modification mark. BRD2 is associated at the strongest level with H4K5acK8ac, followed by H3K27ac and H3K4me3, and at the lowest level with H3K4me1 (**Fig. 3a**). This finding was further verified using an independent analysis where we determined the correlation between BRD2 and H4K5acK8ac or H3K27ac ChIP-seq read intensities within a defined set of active genomic regions. First, we created a reference of 30,243 active genomic regions based on CAGE transcription start sites (unidirectional TSS and bidirectional enhancers), H4K5acK8ac peaks, and H3K27ac peaks. Based on this reference, normalized ChIP-seq read intensities of BRD2 were plotted against those of H4K5acK8ac or H3K27ac (**Fig. 3b**). The levels of H4K5acK8ac correlate well with that of BRD2; weak H4K5acK8ac with weak BRD2 and strong H4K5acK8ac with strong BRD2 (**Fig. 3b**, left), whereas a small number of the active regions is enriched for strong H3K27ac, but contains much weaker BRD2 signals (**Fig. 3b**, right, the top right corner). Thus, BRD2-H4K5acK8ac co-localization shows higher enrichment (Odds Ratio of 62.99) than the BRD2-H3K27ac association (Odds Ratio of 15.32) (**Fig. 3b**). Collectively, our findings uncovered for the first time a strong evidence for the preferential binding of BRD2 to K5/K8-diacetylated histone H4 sites (representing H4-hyperacetylated states) genome-wide.

**Figure 3.**
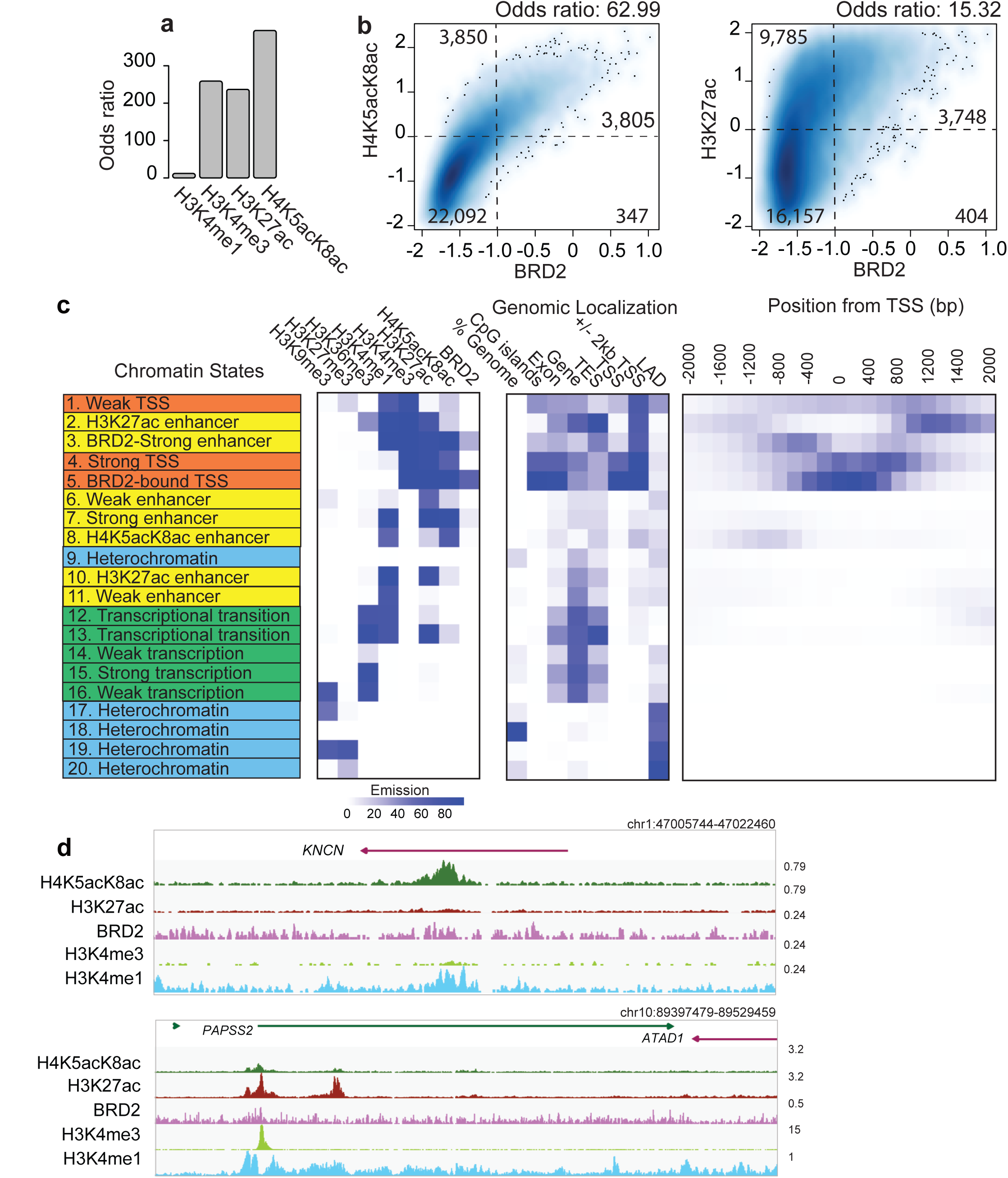
Association of BRD2 with Histone H3 and H4 acetylation and different chromatin states defined by ChromHMM. **(a)** Bar plot showing the genome-wide association of BRD2 binding sites and histone marks (H3K4me1, H3K4me3, H3K27ac and H4K5acK8ac) measured by Odds ratio. The higher Odds ratio the higher association. **(b)** Association between BRD2 binding and H4K5acK8ac (left), H3K27ac (right), shown by a density plot: x-axis: the normalized ChIP-seq signal (log10 RPM + 0.25) of BRD2, y-axis: H4K5acK8ac (left) or H3K27ac (right). The regions were divided into weak and strong signal categories (thresholds: −1 for BRD2; 0 for H4K5acK8ac and H3K27ac) and the numbers represent contingency table between weak and strong categories of the compared marks. The Odd ratios were calculated by Fisher Exact Test using the contingency table as input and represent the co-occurrence/en-richment between strong BRD2 signals and H4K5acK8ac (left) or H3K27ac signals (right). **(c)** 20 ChromHMM chromatin states defined based on the histone marks and BRD2. The 2 left panels represent the predicted chromatin states, shown by different colors (orange for promoter/TSS, yellows for enhancer, green for transcription/elongation, and blue for heterochromatin); the middle panel (genom-ic localization) indicates the genomic distribution of each state; the right panel shows the enrichment of each state within +/-2,000bp of TSS (RefSeq). **(d)** Genome browser views showing examples of H4K5acK8ac-preferential enhancer (upper) and H3K27ac-preferential enhancers (lower).

### Identification of a chromatin state with high H4K5acK8ac, low H3K27ac and low BRD2

We next asked whether there are any regions in the genome that are more highly enriched for H4K5acK8ac than H3K27ac (and *vice versa*), and whether there is any correlation with the BRD2 level. We performed ChromHMM^24^ with 5 active histone marks (H4K5acK8ac, H3K4me1, H3K27ac, H3K4me3, and H3K36me3), 2 inactive histone marks (H3K27me3 and H3K9me3), and BRD2. Based on 20 emission states, we successfully recapitulated the known chromatin state patterns (**Fig. 3c** and **Supplementary Fig. 3a,b**): strong and weak promoters (orange states: 1, 4, 5), enhancer (yellow states 2–3, 6–8, 10–12), transcription (green states: 12–16) as well as heterochromatin (blue states 9, 17–20). The strong promoter states were differentiated into states 4 and 5 by the BRD2 presence (state 5 is BRD2-bound, and state 4 is BRD2-unbound promoters). Of the five strong enhancer states, state 8 ‘H4K5acK8ac enhancer’ is significantly enriched for H4K5acK8ac compared to H3K27ac (**Fig. 3d**, upper), whereas state 10 ‘H3K27ac enhancer’ is more significantly enriched for H3K27ac than H4K5acK8ac (**Fig. 3d**, lower), both of which are not enriched for BRD2 (**Fig. 3c** and **Supplementary Fig. 3c**). Additionally, enhancer state 3 contains both H4K5acK8ac and H3K27ac and is moderately enriched with BRD2 (**Fig. 3c**).

Since the H4K5acK8ac or H3K27ac enhancer states have low BRD2 enrichment, we next investigated the presence of other bromodomain proteins, EP300 (from A459) and BRD4 (from SCLCL H2171) at these states (**Fig. 3c**). Although EP300 and BRD4 are enriched in both enhancer states (**Supplementary Fig. 3e**), BRD4 enrichment is much higher in H4K5acK8ac enhancer state than in H3K27ac enhancer state (**Supplementary Fig. 3e**, left and right), again suggesting preferential binding of BRD4 to H4K5acK8ac. Overall, our analyses suggest that different bromodomain-containing proteins might recognize H4K5acK8ac- or H3K27ac-specific enhancers. Furthermore, although BRD2 has a preferred association to H4K5acK8ac over H3K27ac, it tends to be found at promoters/enhancers that have both H4K5acK8ac and H3K27ac.

### BET inhibition by JQ1 does not affect global K5acK8ac level of histone H4

Since BRD2 exhibits a preference to H4K5acK8ac, we sought to elucidate whether inhibition of BET protein binding by JQ1 affects hyperacetylated histone H4 level. To examine this, H23 cells were treated with 500 nM JQ1 for 24 hours followed by “spike-in” ChIP-seq with either BRD2 or H4K5acK8ac antibody (see **Methods**). Under this condition, BRD2 binding was almost completely depleted at a global level (**Fig. 4a**). Consistent with the previous studies^5,15^, we did not observe any global changes in H3K27ac level (**Supplementary Fig. 4b**). Interestingly, we observed that global H4K5acK8ac levels (including at promoters, enhancers, and super-enhancers) were also not affected by JQ1 (**Fig. 4a**, middle and right, **Fig. 4b,c**). Thus, the inhibition of BET protein binding to chromatin does not significantly change the global hyperacetylation level of histone H4, implying that BET proteins are not required for maintaining or protecting the H4K5acK8ac histone modifications.

**Figure 4.**
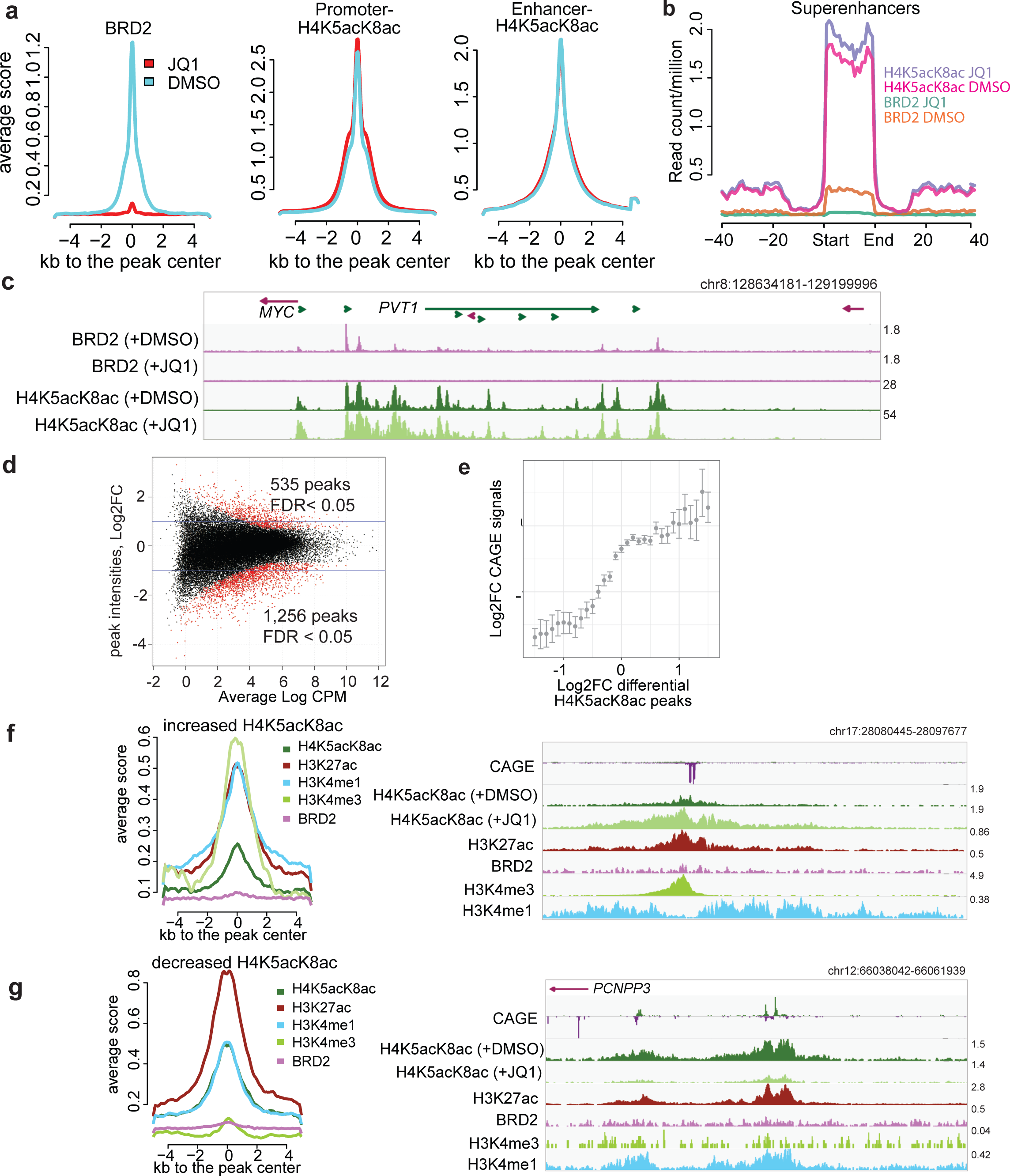
Effects of BET inhibition by JQ1 on BRD2 and H4K5acK8ac sites. **(a)** Treatment with 500 nM JQ1 for 24 hours led to global reduction of BRD2 binding sites (left) but not H4K5acK8ac at promoter (middle) and enhancer (right) regions. Average normalized read intensities of BRD2 and H4K5acK8ac ChIP-seq from JQ1 (red) or DMSO (blue)-treated cells were plotted on the BRD2 or H4K5acK8ac peaks derived from untreated cells, respectively. **(b)** Enrichment of H4K5acK8ac and BRD2 signal in the super-enhancer upon JQ1 treatment. **(c)** Genome browser view of BRD2 binding (top) and H4K5acK8ac signal (bottom) around the MYC-PVT1 loci in the JQ1- and DMSO treated cells. **(d)** MA scatter plot showing the log2 fold changes (Log2FC >1 or >-1, FDR < 0.05) of 535 increased and 1,256 reduced H4K5acK8ac peaks upon JQ1 treatment (y-axis) vs mean intensity (x axis). **(e)** Relationship between changes in the H4K5acK8ac ChIP-seq peaks’ signal (Log2FC, x-axis) and the expression changes of their associated genes (Log2FC, y-axis) upon JQ1 treatment. **(f)** Enrichment of H3K4me3, H3K27ac, H3K4me1 and BRD2 within 5 kb +/-from the summit of peaks with increased H4K5acK8ac signal (n=553) upon JQ1 treatment, shown by normalized average profile. Example of the affected regions is shown on the right. **(g)** Enrichment of promoter and enhancer marks within 5 kb +/-from the summit of peaks with H4K5acK8ac decreased signal (n=1,256) Example of the affected regions is shown on the right.

Despite the lack of global H4K5acK8ac change, we observed a small subpopulation of H4K5acK8ac peaks (~2%; false discovery rate, FDR ≤ 0.05, 2-fold change) to be sensitive to JQ1 treatment (**Fig. 4d**). Similar changes in H3K27ac levels were also observed at these regions (**Supplementary Fig. 4c,d**) and the changes in the H4K5acK8ac levels correlate significantly with the transcriptional activities of the associated promoters (**Fig. 4e**). The increased H4K5acK8ac peaks are associated with promoters, as they are highly enriched for H3K4me3 and to a lesser degree for H3K4me1 (**Fig. 4f**), while the decreased peaks are associated with transcribed enhancers, indicated by CAGE peaks and the high H3K4me1/H3K4me3 ratio (**Fig. 4g**). Interestingly, the H4K5acK8ac-specific regions that are affected by JQ1 are not bound by BRD2, although we cannot exclude BRD4/BRD3 enrichment in these regions (**Fig. 4f,g**).

### JQ1 treatment induces apoptosis in a *MYC*-independent manner in H23 cells

We next asked how the H4K5acK8ac/H3K27ac ratios are associated with transcriptional response to JQ1. We observed that active promoters with higher H4K5acK8ac level tend to be more downregulated upon JQ1 treatment (**Fig. 5a**, bottom right corner), whereas active promoters with similar levels of H4K5acK8ac/H3K27ac or high H3K27ac show both directions (*i.e.,* upregulated and downregulated), suggesting different regulatory mechanism are acting on these regions.

**Figure 5.**
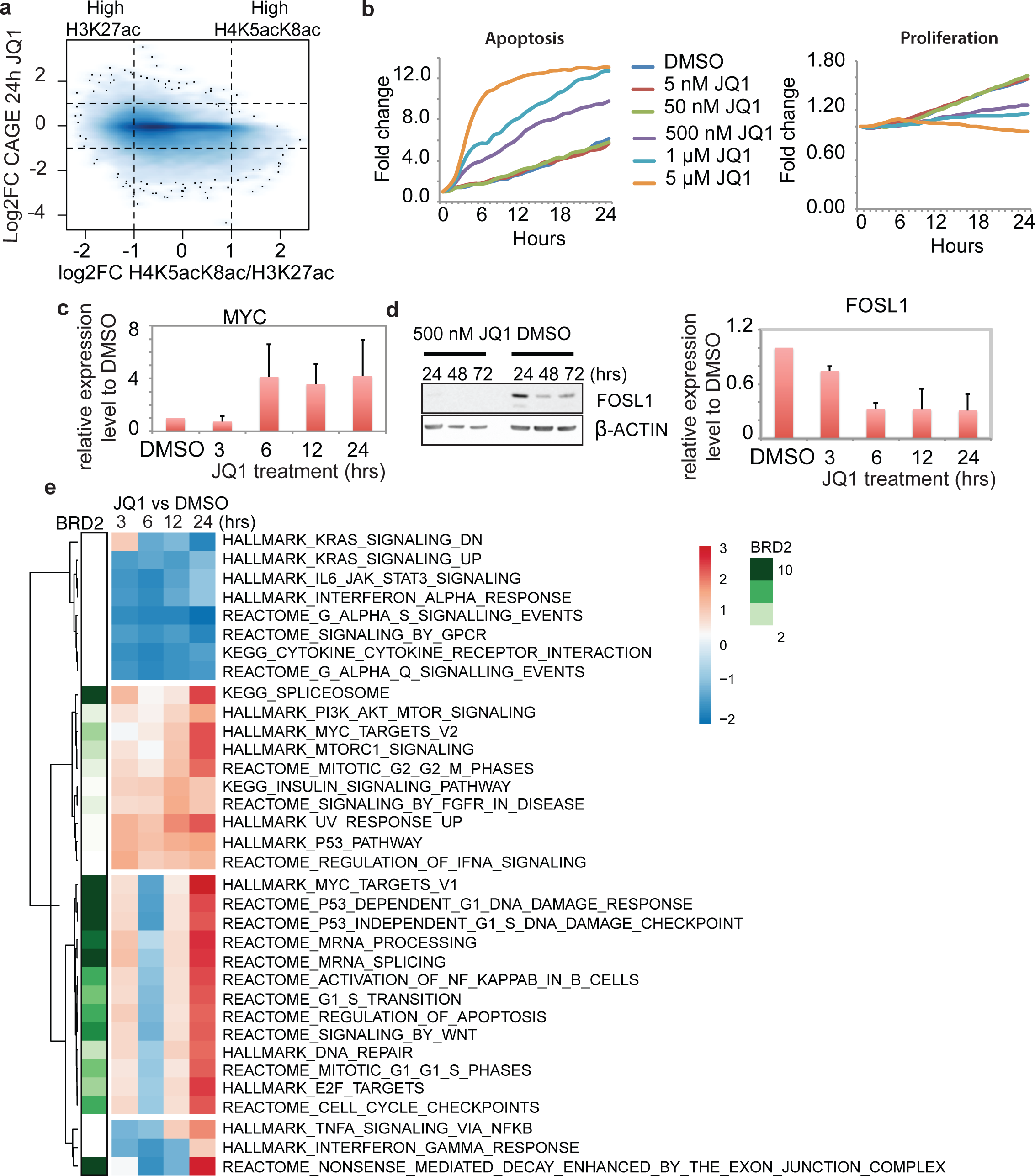
Effects of BET inhibition by JQ1 the transcriptional regulation in H23 cells. **(a)** Density plot of H4K5acK8ac/H3K27ac ratio (x-axis) versus the transcriptional response (log2 fold change) to JQ1 treatment (24 hours) measured by CAGE (y-axis). The values were calculated within 1 kb window of 30,243 active centers. Noteworthy, a fraction of H4K5acK8ac-enriched sites are downregulated after JQ1 treatment, but virtually no H4K5acK8ac-enriched sites are upregulated. **(b)** Effects of different concentrations of JQ1 (5 nM, 50 nM, 500 nM, 1 µM, and 5 µM) on the cell apoptosis (left) and proliferation (right) during 24 hours (x-axis). The y-axis reflects the fold change of the number of the apoptotic (left) or proliferating (right cells at the indicated hours (1-24 hours) compared to 0h. **(c)** MYC expression upon JQ1 treatment at different time points. The gene expression level was determined by quantitative PCR from the JQ1- and DMSO-treated cells and normalized against GAPDH. **(d)** FOSL1 expression upon JQ1 treatment at 24-72 hours. Left: FOSL1 expression upon 500 nM JQ1 (left) and DMSO (right) treatment, as examined by Western blot. Right: FOSL1 gene expression level upon JQ1 treatment at 3, 6, 12, 24 hours, as validated by qPCR, the expression level was normalized against DMSO and GAPDH. **(e)** Heatmap of the pathways after JQ1 treatment and association of BRD2 with the upregulated pathways. The clustering was performed based on the normalized enrichment score (NES) of the enriched pathways (p-value ≤ 0.01, FDR < 0.25) and visualized as a color-coded matrix. Color density indicates the enrichment; the left panel shows association of BRD2 binding sites with each pathway.

Differential expression analysis between JQ1-treated cells and DMSO-treated (vehicle control) cells at 3, 6, 12 and 24 hours showed that more genes were downregulated than upregulated (FDR < 0.05 and fold change ≥ 2, **Supplementary Fig. 5a** and **Supplementary Table 5**). We successfully validated 20 differentially expressed genes by qPCR (**Supplementary Fig. 5b** and **Supplementary Table 9**). Consistent with other studies^5,15^, *BRD2* and *BRD4* expression levels were not reduced by JQ1 treatment (confirmed by qPCR), and similar to other NSCLC cell lines^25^, anti-apoptotic oncogenes *CFLAR* (*FLIP*) and *BCL2* were markedly downregulated after 3 and 24 hours of JQ1 treatment, respectively (**Supplementary Fig. 5c**). *HOXC* genes (important for anti-apoptotic pathway^26^) were downregulated at the early time point (**Supplementary Fig. 5d**) in line with the number of H23 cells undergoing apoptosis and cell proliferation rate (**Fig. 5b**). The main components of canonical Wnt/β-catenin signaling are reported to be modulated by JQ1^5,27,28^, and in H23 cells, *WNT2*, *WNT5A*, and the mediator of Wnt signaling (*LEF1* and *TCF* genes) were downregulated, while β-catenin encoded by *CTNNB1* was only slightly affected (**Supplementary Fig. 5e**). *MYC* is a Wnt signaling target and a key transcription factor (TF) that is downregulated by JQ1 in MM.1S, leukemia, and breast cancer cells ^5,15,29^ and is believed to mediate JQ1’s anti-oncogenic effect. Here we observed the upregulation of *MYC* expression after JQ1 treatment in H23 cells (**Fig. 5c** and **Supplementary Table 4**). This is in agreement with results from another JQ1-sensitive NSCLC cell line^10^, where the effect of JQ1 on the cell growth is thought to be mediated by *FOSL1*, and not *MYC*^10^. Notably, we also observed the downregulation of *FOSL1* upon JQ1, as confirmed by Western blot and qPCR (**Fig. 5d**).

### Potential BRD2-coupled transcription regulatory pathways

To elucidate the regulatory pathways affected by BET inhibition, we carried out pathway analysis upon JQ1 treatment at each time point and determined BRD2 enrichment in these pathways. Specifically, we performed a clustering on the normalized enrichment score (NES) of the enriched pathways (p-value ≤ 0.01, FDR < 0.25) (**Fig. 5e**, the complete list of the enriched pathways is available in the **Supplementary Table 6**). Upregulated pathways, such as cell cycle regulation, Wnt/β-catenin signaling target genes, apoptosis pathway, E2F-target genes, RNA processing-related genes, and MYC target genes, were highly enriched for BRD2 binding (**Fig. 5e** and **Supplementary Fig. 5d,e**). On the other hand, downregulated pathways, such as cytokine–cytokine receptor pathway, G protein signaling, and *KRAS* target genes, were much less enriched for BRD2 binding (**Fig. 5e** and **Supplementary Fig. 5d,e**). Differential expression analysis of the BRD2-bound genes also shows similar sets of genes are enriched (FDR < 0.05, log_2_ fold change ≤ −1; **Supplementary Table 7**), including upregulation of *MYC*. Furthermore, pathway analysis using multiple functionally validated gene sets confirmed the enrichment of MYC and E2F target gene sets in the upregulated genes (NES ≥ 1.5, FDR ≤ 0.01) (**Supplementary Fig. 5f**).

### Identifying TF candidates of BRD2-dependent and independent regulatory pathways

Since JQ1 treatment shows diverse effects (both upregulation and downregulation) on BRD2-bound genes, we hypothesized that different TFs might be involved in the transcriptional regulation in response to JQ1. We first performed Motif Activity Response Analysis (MARA)^30^ (Z-value ≥ 2.0) using CAGE data and identified several candidate TFs that are responsible for up- or downregulation of genes upon JQ1 treatment (Z-value ≥ 2.0) (**Fig. 6a** and **Supplementary Table 8**). Among them are *E2F3* and *YY1* that show increased motif activity upon JQ1 treatment, and *ZNF281*, *LEF1*, *FOSL1* that show decreased motif activity (**Fig. 6a**). To further refine TF candidates into BRD2-dependent or independent pathways, we performed motif search analysis by HOMER^31^ using BRD2 binding sites of upregulated and downregulated promoters. The downregulated promoters were enriched in ZNF281 binding motifs (**Fig. 6b**, left), while E2F4 and YY1-binding motifs were enriched in the upregulated promoters (**Fig. 6b**, right). Since *LEF1* and *FOSL1* were not identified in the downregulated genes, they are likely to be involved in the non-BRD2 regulatory pathway. To investigate this, we performed HOMER^31^ motif search and found that LEF1 and FOSL1 binding motifs were enriched within H4K5acK8ac peaks without BRD2 (**Fig. 6c**, right). On the other hand, E2F, YY1, and ELK1 motifs were enriched in H4K5acK8ac peaks with BRD2 binding (**Fig. 6c**, left). It is of note that previous studies revealed interaction of BRD2 with CTCF^32,33^. Our HOMER motif enrichment analysis also shows higher enrichment of CTCF motif in the downregulated genes (**Supplementary Fig. 6a**) and the CTCF binding sites (from A549 cells) are more enriched in the BRD2 peaks associated with the downregulated genes than upregulated genes (**Supplementary Fig. 6b**). Taken together, our findings provide a possible model where BRD2 recruits TFs such as YY1, E2F4, CTCF, and ZFN281 and regulate these genes, and other TFs such as LEF1 and FOSL1 are recruited to non-BRD2 BET proteins.

**Figure 6.**
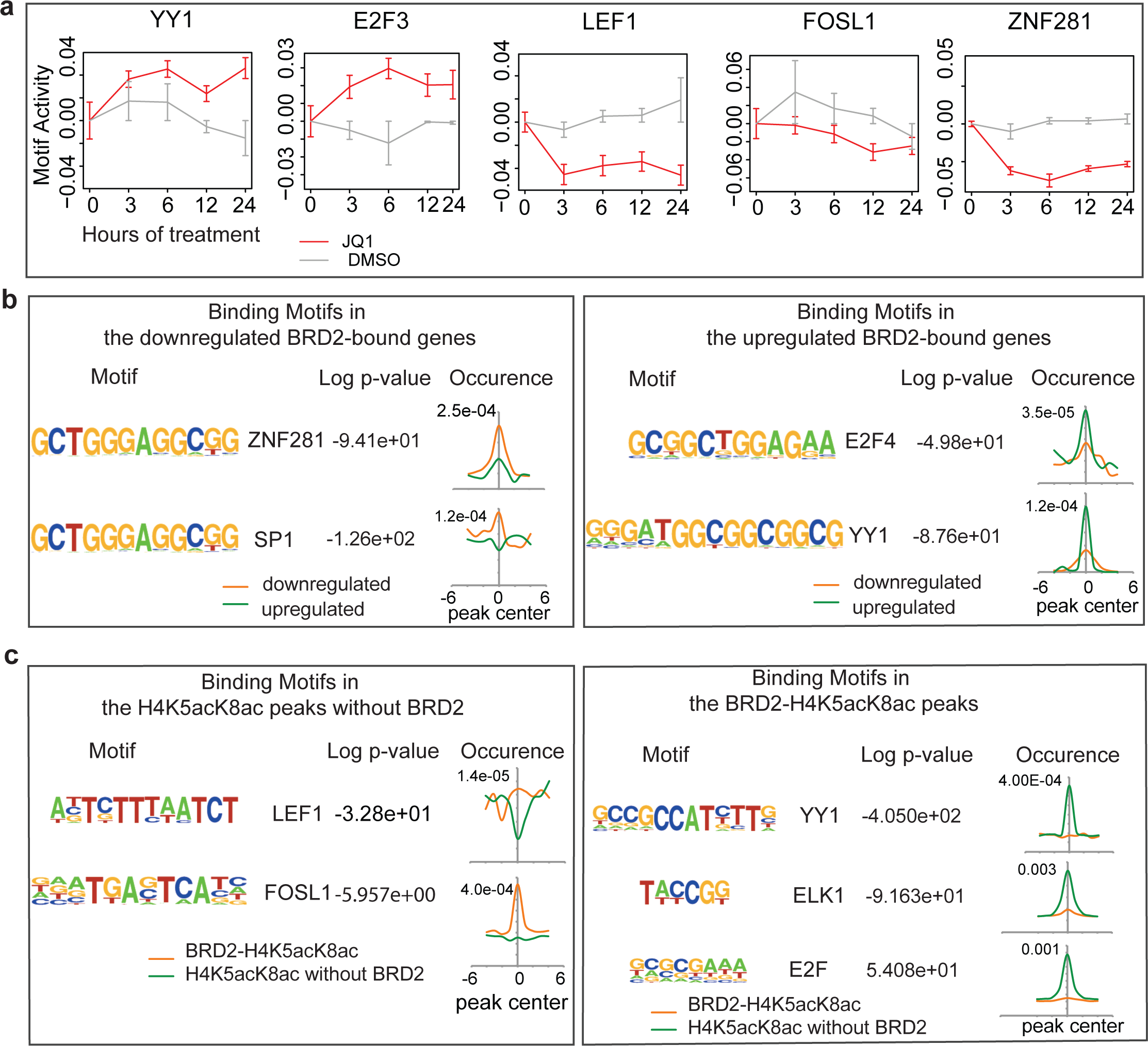
Association of BRD2 with different transcription factors. **(a)** The changes of motif activity of YY1, E2F3, LEF1, FOSL1, and ZNF281 transcription factors estimated by Motif Activity Response Analysis (MARA). Red and gray lines indicate the motif activities in the JQ1- and DMSO-treated cells, respectively. The motif activities reflect the expression of genes regulated by the transcription factors. **(b)** Transcription factor-binding motifs within the BRD2 peaks that overlap the downregulated (right) or the upregulated genes (left). The motifs were found de novo by HOMER and then linked to the best matching known motifs. The occurrence panels represent the enrichment of the motif within 6 kb +/-from the peak summit in BRD2 peaks at the downregulated (orange) and upregulated (green) genes. **(c)** LEF1 and FOSL1-binding motifs enriched in H4K5acK8ac peaks without BRD2 (right) and E2F, ELK1, YY1 binding motifs enriched in the BRD2-H4K5acK8ac peaks (left).

## DISCUSSION

Antibody quality in terms of target specificity and reproducibility is one of emerging issues in the biological sciences^34,35^. In this study, we generated monoclonal antibody that specifically detects di-acetylation at K5 and K8 of histone H4, a hallmark of H4-hyperacetylated chromatin states *in vivo*. Peptide array and crystal structure analyses confirmed its exclusive binding to H4K5acK8ac. By applying the structurally validated antibody to ChIP-seq, we demonstrate that BRD2 preferentially associates with H4K5acK8ac genome-wide. We showed that the reduced binding of BRD2 by JQ1 did not affect the global H4K5acK8ac level at the BRD2 sites. We further identified active enhancer states containing significantly different enrichment between H4K5acK8ac and H3K27ac. By separately analyzing the differentially expressed regions that are bound or not bound by BRD2 prior to the JQ1 treatment, we identified several TF candidates that may be involved in the regulation of BRD2 and non-BRD2 target genes.

The finding that H4K5acK8ac levels at BRD2 binding sites were not affected despite complete reduction of BRD2 after JQ1 was surprising. In LPS-activated macrophages, BET inhibition reduced the levels of H3ac, H4K5ac, H4K8ac, H4K12ac and total H4ac on promoter of the induced genes^36^. This reduction is presumably due to increased accessibility of HDAC to the exposed BET sites by preventing the formation of multi-molecular complexes containing HATs and other transcription machineries^36^, as shown by the reduced level of positive transcription elongation factor b (P-TEFb/*CDK9*) and RNA polymerase II on the affected regions^36^. In H23 cells, however, a subset of BRD2-bound genes involved in RNA processing such as *POL2RA* and *CDK9* was in fact upregulated, and the transcription levels of HATs, such as *KAT2A* and *EP300*, were not affected by JQ1. This indicates that HAT-containing transcription complexes can still be recruited upon JQ1 treatment, and H4-hyperacetylated states could be maintained through the HAT activities of these complexes even if BET proteins are off the chromatin.

Several JQ1 studies have been conducted in NSCLC cell lines^10,19^, however, our study delivers more comprehensive transcriptome data by using CAGE, with which we identified downregulation of cytokine–cytokine receptor interaction pathways and several regulators of tumor-associated inflammation, such as IL6, which were not reported before. These findings are consistent with the previous reports that JQ1 inhibits inflammation in pancreatic ductal adenocarcinoma^37^ and a murine periodontitis model^38^. Thus, our study implicates that JQ1 might also reduce inflammation in NSCLC cells.

Although *MYC* is downregulated in many cell types upon JQ1 treatment and thought to be the key mediator of JQ1 effects, our study together with the previous study^10^ suggests that H23 cells exhibit MYC-independent response to JQ1. Supporting the previous study that proposed FOSL1-dependent mechanism^10^ in response to JQ1, our study suggests FOSL1 and LEF1 might regulate transcription of non-BRD2 target genes (quite possibly through BRD3 and/or BRD4). This result is consistent with the previous report that LEF1 acts as a coactivator in the presence of EP300^39^ and induces lung adenocarcinoma metastasis^40^, while FOSL1 controls gene expression program mediating metastasis^41^. Since *FOSL1* and *LEF1* expression levels were reduced by JQ1 treatment, downregulation of their downstream target genes such as *IL6*, *FOS*, and *JUN* may lead to reduction of metastasis.

Furthermore, by integrating the BRD2 ChIP-seq and CAGE data upon JQ1 treatment, we identified YY1/E2F4 transcription factors might be responsible for upregulating and ZNF281 for downregulating the genes upon JQ1 treatment. YY1 and E2F4 are associated with apoptosis and proliferation in tumor cells, and can act as a transcription activator or a repressor (directly or indirectly) via cofactor recruitment ^42–45^. In our study, the expression of YY1 and E2F4 were not affected by JQ1. Thus, they may still bind to their target genes and recruit other coactivators that lead to up-regulation of their target genes. Since ZNF281 is involved in the control of epithelial–mesenchymal transition (EMT) and promotes metastatis^46^, anti-oncogenic effects of JQ1 on H23 cells may partly be due to downregulation of ZNF281, which may lead to apoptosis. Further analyses, such as identification of their target genes by ChIP-seq, knockdown experiments of the TFs and immunoprecipitation of BRD2 protein complex would elucidate the roles of these TFs.

Collectively, we shed lights on the mechanism of BET inhibition effects at the H4-hyperacetylated chromatin and transcriptome levels in H23 cells. The complexity of BET inhibition effects becomes increasingly evident, and it can be explained by future comparative analyses among BET family members, cell types, and epigenetics-regulating agents. In particular, our study reveals that BET inhibitors such as JQ1 may only be effective temporarily as anti-cancer progression, since they may not have the potency to change the epigenomic state of H4 hyperacetylation by monotherapy. It would, therefore, be interesting to investigate the synergistic effects of BET and HAT inhibitors on chromatic histone acetylations, especially on H4 hyperacetylation.

## METHODS

Methods and associated references are available in the supplementary file.

### Accession codes

The atomic coordinates and structure factors have been deposited in the Protein Data Bank (PDB) under the following accession codes: The Fab fragment of 2A7D9 in complex with the H4(1–12)K5acK8ac peptide: 5YE3; The Fab fragment of 1A9D7 in complex with the H4(1–12)K5acK8ac peptide: 5YE4 (**Supplementary Table 2**). The ChIP-seq data used in this study have been deposited in the Gene Expression Omnibus (GEO) under accession code GSE104481.

## ACKNOWLEDGEMENTS

We acknowledge Dr. Naohito Nozaki (Mab Institute Inc.) for his help in developing anti H4K5acK8ac antibodies; Dr. Atsushi Miyawaki (RIKEN Brain Science Institute) for providing us with the plasmid containing Venus; Dr. Hiroyuki Miyoshi (Keio University) for developing it; the RIKEN BioResource Center (RIKEN BRC) for distributing it; and the BL26B2 beamline staff at the SPring-8 for their support in data collection; Drs. Kazuhide Watanabe and Supat Thongjuea for the discussion; and Ms. Haruka Yabukami for technical supports. This work was supported by research grants from the Japanese Ministry of Education, Culture, Sports, Science and Technology (MEXT) to the RIKEN Center for Life Science Technologies (CLST); the Center Director’s Strategic Fund of CLST; the PRESTO program of the Japan Science and Technology Agency (JST); and Grant-in-Aid for Scientific Research (B) (16H05089) from the Japan Society for the Promotion of Science (JSPS). B.K. was supported by Foreign Postdoctoral Researcher (FPR) program from RIKEN, Japan.

## AUTHOR CONTRIBUTIONS

L.H. performed most of the experiments; B.K., L.H., C.C.H., and M.L. performed the bioinformatics analyses. M.W., K.S., S.Y., and T.U. generated the antigens and antibodies. T.M. and T.I. performed crystal structure analysis. P.J. performed validation qPCR experiments. Y.S. and H.K. performed immunostaining analysis. L.H., B.K., A.M., and T.U. designed the experiments, interpreted the results and wrote the manuscript. A.M. and T.U. supervised the project. All authors commented on the manuscript.

